# “Identification of microRNAs regulated by E2F transcription factors in human pluripotent stem cells”

**DOI:** 10.1101/2024.02.28.582539

**Authors:** María Soledad Rodríguez-Varela, Mercedes Florencia Vautier, Sofía Mucci, Luciana Isaja, Elmer Fernández, Gustavo Emilio Sevlever, María Elida Scassa, Leonardo Romorini

**Affiliations:** Laboratorios de Investigación Aplicada en Neurociencias (LIAN), Instituto de Neurociencias (INEU-CONICET), Fundación para la Lucha contra las Enfermedades Neurológicas de la Infancia (Fleni), Ruta 9, Km 52.5, Belén de Escobar, Provincia de Buenos Aires, B1625XAF, Argentina; DataLab, Fundación para el Progreso de la Medicina, CONiCET, Córdoba, Argentina; Facultad de Ciencias Exactas, Físicas y Naturales, Universidad Nacional de Córdoba, Córdoba, Argentina

**Keywords:** human pluripotent stem cells, E2Fs, microRNAs

## Abstract

Human pluripotent stem cells (hPSCs), which include embryonic and induced pluripotent stem cells (hESCs and hiPSCs, respectively), have an unusual cell cycle structure which consists of a short G1 phase and the absence of the G1/S checkpoint regulation. E2F transcription factors (E2Fs) play an important role in the G1/S transition. G1 duration contributes to hPSC fate determination, and microRNAs (miRNAs) play critical roles in this commitment. As little is known about the interplay between E2Fs and miRNAs in hPSCs, we aimed to identify miRNAs that are regulated by E2Fs in these cells. We first found that mRNA expression levels of canonical E2F repressors were more expressed than most E2F activators in G1-arrested hPSCs. Moreover, we observed higher mRNA and protein expression levels of canonical *E2F2*, *E2F3A,* and *E2F5* in G1 synchronized hPSCs compared to human fibroblasts (HF). However, *E2F1* and *E2F4* protein expression levels were higher in HF. We next found that E2F inhibition with HLM006474 induced an increase in the G1 cell population without affecting hPSC viability, concomitantly with a decrease in *OCT-4* mRNA levels and the percentage of OCT-4^+^ hPSCs. Next, by RNA-seq analysis we found 52 differentially expressed (DEGs) miRNAs in HLM006474-treated hESCs. RT-qPCR validation of some of the DEGs let us conclude that miR-19a-3p, miR-19b-3p, miR-4454, miR-1260a, miR-1260b, miR-454-3p and miR-301a-3p are regulated by E2Fs in hPSCs. Interestingly, gene target and ontology analysis of these miRNAs revealed a possible implication in proliferation and cell cycle regulation, development, and neural differentiation.

## INTRODUCTION

Human embryonic and induced pluripotent stem cells (hESCs and hiPSCs, respectively), are self-renewing pluripotent stem cells (hPSCs) that can differentiate into all somatic and germ cell lineages [1–3]. The ability of hPSCs to maintain their self-renewal and pluripotency is associated with their capacity to remain in a proliferative condition, which requires a unique transcriptional profile [4]. The hPSC cell cycle is unique, characterized by cells that are mainly transiting the S phase (65% of the time), a shortened G1 phase (15% of the time), and the absence of G1/S checkpoints [5]. Although there are exceptions, hPSCs divide quickly with generation times between 8-16 hours [6–8]. When hPSCs differentiate, cells accumulate in the G1 phase and show an elongated cell cycle of more than 16 hours, as seen in somatic cells. The shortened G1 phase increases the rate of cell division since the duration of the S phase is like other cell types [9, 10]. Despite great progress in understanding the molecular mechanisms involved in regulating the hPSC cell cycle, several aspects remain unknown.

The progression through the cell cycle is accompanied by an increase in the expression of a set of specific genes or regulatory RNAs during one phase of the cell cycle and a decrease in the same set of genes during a later phase of the cell cycle [11]. Particularly, microRNAs (miRNAs), which are non-coding RNA molecules of approximately 18-24 nucleotides that regulate gene expression by interacting with specific sequences present in mRNAs, are implicated in fine-tuning numerous biological processes such as cell cycle progression, proliferation, and differentiation [12]. Interestingly, some miRNAs promote hPSC proliferation [13], and some are regulated in a cell cycle-dependent manner [14].

It has become evident that E2F transcription factors play a crucial role in regulating stem cell fate control across various lineages, and in mammals, this role extends to pluripotent stem cells [15]. E2F transcription factors are critical for the temporal expression of cell cycle oscillating genes and non-coding RNAs, like miRNAs [11]. In quiescent cells, growth factors activate G1 phase Cyclin-Cyclin dependent kinase (CDK) complexes, which in turn hyperphosphorylate members of the Retinoblastoma (pRB) family of proteins, pRB, p107, and p130 that are bound to E2Fs. This leads to a conformational change of pRB, thus releasing E2F factors from the complex [16]. To date, eigth E2F genes, giving rise to ten distinct proteins, have been identified in mammals, which are divided into activators (E2F1–3A) and repressors (E2F3B, E2F4– 8). E2F1–6 are classic or canonical E2Fs, with a DNA-binding domain, and are required to form heterodimers with the DP1/DP2 proteins before they can bind to the promoters of the target genes and activate or repress their expression. E2F7 and E2F8 are considered atypical E2Fs. They have two DNA-binding domains and can repress target genes independently of heterodimerization with DP proteins [17]. The accumulation of free E2F activators then leads to an increase/decrease in the expression of their targets. The E2F activators primarily bind to promoters of genes or non-coding RNAs that are important for S phase progression leading to the initiation of DNA replication.

hPSCs lack the typical pRB/E2F-mediated transition in the late G1 phase, which normally sustains the commitment of entry to the S phase at the restriction point [18]. The absence of this pRB/E2F switch in pluripotent cells predicts a mechanism that prevails over the entry of hPSCs into the S phase without the requirement of supplementary exogenous factors. In this sense, Becker *et al*. demonstrated that hESCs are already involved in mitosis upon entry to the next S phase without the presence of external stimuli [19]. Due to the high activity of Cyclins/CDKs complexes observed in hPSCs, it would be expected that members of the pRB family are not able to bind to E2Fs, thus allowing E2F1–3 to activate their target genes/non-coding RNAs and thus leaving E2F4–5 without binding to DNA.

Since miRNAs are transcribed by RNA polymerase II, they can be regulated by transcription factors. This fact predicts that regulatory loops exist between genes coding for transcription factors and genes coding for miRNAs. To our knowledge, little is known about the interplay between E2Fs and miRNAs in hPSCs. So, herein, we aimed to elucidate whether E2Fs regulate miRNAs in hESCs and hiPSCs. We determined higher mRNA expression levels of canonical *E2Fs* in hPSCs arrested in early G1 with PD0332991 compared to somatic cells (human fibroblast, HF). We observed that *E2F2*, *E2F3a,* and *E2F5* are more abundant at the protein level in hiPSCs than in FH. On the contrary, we found higher protein expression levels of *E2F1,* and *E2F4* in HF compared to early G1 synchronized hPSCs. Moreover, interestingly we found that E2F repressors (*E2F3B*, *E2F4* and *E2F5*) mRNAs are more expressed than most activator E2Fs in G1-arrested hPSCs, except for E2F2.

Then, we treated hPSCs with the general E2Fs inhibitor (pan-E2F inhibitor) HLM006474 (20 ⎧M) for 24 hours and observed an increase in G1 cell population, a decrease in *OCT-4* mRNA expression levels and OCT-4^+^ cells and no effect in hPSC viability. Furthermore, we found that E2F inhibition decreased *CYCLIN A2* mRNA expression levels in H9 hESCs and increased *E2F1* mRNA expression levels in FN21 hiPSCs. These genes were chosen as they were reported to be regulated by E2F transcription factors in other biological models. These results highlight the connection between the cell cycle machinery that regulates the G1/S phase transition and the pluripotency network.

Finally, we performed RNA-seq analysis of small RNAs and found 52 differentially expressed miRNAs in H9 hESCs upon HLM006474 inhibition. Within these, some were already associated with E2Fs while others had no reported relationship with these factors or the hPSC-cell cycle. After validating the changes in the expression levels of some of these candidates by specific RT-qPCR, we concluded that E2Fs would be transcriptionally regulating miR-19a-3p, miR-19b-3p, miR-4454, miR-1260a, miR-1260b, miR-454-3p, and miR-301a-3p not only in hESCs but also in hiPSCs. Interestingly, gene target and ontology analysis of these miRNAs revealed a possible implication in development and differentiation processes.

## MATERIALS AND METHODS

### Cell lines and culture

hESCs line WA09 (H9) [2] was purchased from WiCell Research Institute (http://www.wicell.org) and hiPSCs line FN2.1 was derived from human foreskin fibroblasts (HF) [20]. Ethical approval was provided by the local Ethics Committee (Comité de Ética en Investigaciones Biomédicas del Instituto Fleni) and the donor give a written informed consent before foreskin fibroblast isolation. hPSCs were cultured on Geltrex (Thermo Scientific) coated dishes in combination with fully defined mTeSR1 medium (Stem Cell Technologies). HF were prepared as primary cultures from freshly obtained human foreskins as soon as possible after surgery as previously described [8]. All cell lines were free of *Mycoplasma sp.* infection, which was tested as previously described [21].

### RNA isolation and RT-qPCR

Total RNA was extracted using TRIzol reagent (Thermo Scientific) according to the manufacturer’s instructions. 500 ng of total RNA was used for cDNA synthesis with 15 mM of random hexamers and MMLV reverse transcriptase (Promega), according to the manufacturer’s instructions. For miRNA reverse transcription, cDNA was generated using SuperScript™ II Reverse Transcriptase (Thermo Scientific) and miRNA-specific stem-loop primers as previously described [14]. For real-time PCR, cDNA samples were diluted 5-folds and PCR was performed with StepOne Plus Real-Time PCR System (PE Applied Biosystems). The FastStart Universal SYBR Green Master (Roche) was used for all reactions, following the manufactureŕs instructions. For information about primer sequences please see Table S1 and Table S2 in Supplementary information.

### Protein analysis

Protein expression levels were analyzed as previously described [22]. Total proteins extraction from hPSCs was performed in ice-cold RIPA protein extraction buffer (Sigma) supplemented with protease inhibitors (Protease inhibitor cocktail set I, Calbiochem).

Bicinchoninic Acid Protein Assay (Pierce) was used for protein concentration determination. Equal amounts of protein were electrophoresed on a 12% SDS– polyacrylamide gel and transferred to PVDF membranes (Millipore). Blots were blocked for 1 hour at RT (room temperature) in TBS (20 mM Tris–HCl, pH 7.5, 500 mM NaCl) containing low-fat powdered milk (5%) and Tween 20 (0.1%). Incubations with primary antibodies were performed ON (overnight) at 4 °C in blocking buffer (3% skim milk, 0.1% Tween, in Tris-buffered saline). Membranes were then incubated with the corresponding counter-antibody and the proteins were revealed by enhanced chemiluminescence detection (SuperSignal West Femto System, Thermo Scientific). For information about the antibodies used please see Supplementary Table S3. ImageJ 1.34 s software (https://imagej.nih.gov/ij/) was used for densitometric protein level analysis.

### Quantification of OCT-4 positive cells

The percentages of Octamer-binding transcription factor 4 (OCT-4) positive hESCs and hiPSCs were calculated by flow cytometry. Briefly, hPSCs were dissociated by incubation with Accutase, and cell staining was carried out on cells fixed with 4% paraformaldehyde in PBS with 0.5% BSA and 0.5% Saponin (Sigma). Anti-OCT-4 (1:1000) primary antibody from Santa Cruz Biotechnology (sc-5279) and Alexa Fluor 488 goat anti-mouse IgG (1:400) from Invitrogen (A11001) were used. A BD Accuri cytometer was used for flow cytometry analyses. Data were analyzed with FlowJo software.

### Cell viability assay

hPSCs were plated onto Geltrex-coated 96-well dishes at densities between 3.33□×□10^4^–1□×□10^5^ cells/cm^2^ and grown until confluence as previously described [23, 24]. After treatments, 50 μg/well of activated 2,3-bis (2-methoxy-4-nitro-5-sulfophenyl)-5 [(phenylamino) carbonyl]-2 H-tetrazolium hydroxide (XTT) in PBS containing 0.3 μg/well of N-methyl dibenzopyrazine methyl sulfate (PMS) were added (final volume 100 μl) and incubated for 1-2 hours at 37°C. Cellular metabolic activity was determined spectrophotometrically at 450 nm.

### Trypan blue staining

For the Trypan blue exclusion assay, hPSCs were seeded on Geltrex-coated 6-well tissue culture plates at a density of 1□×□10^5^ cells/cm^2^. After treatments, adherent and detached cells were collected and stained with 0.4% Trypan blue solution (final concentration 0.08%) for 5 minutes at room temperature as previously described [23]. Cells were counted in a hemocytometer chamber. Percentages of surviving cells (unstained) were calculated as the total number of live cells divided by the total number of cells (stained) and multiplied by 100.

### Flow cytometric analysis of cell viability by Propidium Iodide (PI) staining

Single-cell suspensions were obtained with Accutase (37°C for 7 minutes). hPSCs were then centrifuged at 200 x g for 5 minutes and resuspended up to 1×10^6^ cells/ml in FACS Buffer (2.5 mM CaCl_2_, 140 mM NaCl and 10 mM HEPES pH 7.4). Next, 100 µl of cellular suspension was incubated with 5 µl of PI (50 ⎧g/ml) in PBS for 5 minutes in the dark. Finally, 400 µl of FACS Buffer was added to each tube and cells were immediately analyzed by flow cytometry [23]. Data were acquired on a BD Accuri C6 flow cytometer and analyzed using BD Accuri C6 software.

### Flow cytometric analysis of bromodeoxyuridine (BrdU) incorporation and cell cycle distribution

The distribution of cell populations throughout the cell cycle and the fraction of cells capable of incorporating BrdU were determined using the BrdU Flow Kit (BD Biosciences). After treatments cells were incubated with BrdU (10 ⎧M) for 30 min. Cultures were processed as per the manufacturer’s instructions. Fluorescence intensity was quantified on a BD Accuri C6 flow cytometer. Data were analyzed using BD AccuriC6 software.

### Flow cytometric analysis of cell cycle by propidium iodide (PI) DNA staining

Single-cell suspensions were obtained with Accutase 1x treatment (37°C for 7 minutes). To analyze DNA content, cells were fixed with ethanol 70%, re-hydrated with PBS 3% FBS, left for 40 minutes at 4°C, then incubated for 30 minutes at 37°C with RNAse A (100 ⎧g/ml) (Invitrogen), and then with Propidium Iodide (PI) (40 ⎧g/ml) (Sigma-Aldrich) for 5 minutes. Cells were immediately analyzed by flow cytometry. Data were acquired on a BD Accuri C6 flow cytometer (BD Biosciences). The percentage of cells in each cell cycle phase was calculated by the FlowJo v10.0.7’s univariate platform which assumes Gaussian distributions of the 2N and 4N (formerly G0/G1 and G2/M, respectively) populations, then uses a subtractive function to identify the S-phase population. The cells within one standard deviation of the 2N and 4N medians are subtracted from the data. The remaining cells (S-phase) fit with a polynomial function that is convoluted with the Gaussian distributions of the 2N and 4N populations to form the complete model.

### Small RNA-Sequencing sample preparation

The RNA was extracted with mirVana^TM^ miRNA Isolation Kit (Ambion, Thermo Fisher Scientific) following the manufacturer’s protocols. We performed a High Throughput Small RNA-Sequencing of total RNA Samples of H9 hESCs control and H9 hESCs treated with the E2Fs pan-inhibitor HLM006474 (iE2Fs; 20 µM for 24 hours). Small RNA libraries were prepared using the TruSeq small RNA Library Prep Kit with adaptors that are also designed to capture small RNAs. Cleanup and size selection were made by gel electrophoresis following the manufacturer’s protocols. The sequencing and initial bioinformatic analysis was carried out in the sequencing facilities of High Throughput Sequencing of the company Macrogen, South Korea (http://foreign.macrogen.co.kr) using an Illumina HiSeq 2500, in one lane and 50bp single-end read sequencing. Each sample was analyzed in triplicate with 21666667 readings per sample. Sequencing datasets are deposited in GEO (GSE253898).

### Data processing and bioinformatic analysis

After sequencing, the raw sequence reads were filtered based on quality and the adapter sequences were also trimmed off the raw sequence reads. The trimmed reads were then aligned and mapped to the human genome reference (GRCh38) with the miRDeep2 mapper and to miRbase (v21) with the miRDeep2 module which allowed us to identify known and novel microRNAs. The read counts for each miRNA were extracted from mapped miRNAs to report the abundance of each miRNA. Bioinformatic analysis was performed in R. Heat maps, box plots, MA plots and principal component analysis (PCA) were plotted using the ggplot2 package. Differential expression analysis was conducted with the Bioconductor EDGER and limma packages using raw data as inputs.

### Gene target analysis

Gene target analysis was done with the miRWalk 3.0 database. Briefly, we selected the first 10 target genes of the 52 differentially expressed miRNAs that have a p-binding ≥ 0.8 and those genes target that the respective miRNA binds to the 3′UTR region of them. Besides, from those genes, we chose the ones that were present in the three or most of the databases that the miRWalk 3.0 software considers (Targetscan, Mirdb, _MiRTarBase). The_ *p*-binding is the probability that a candidate target site is a real target site. For the gene target list please see Table S4 in the Supplementary information.

### Gene ontology analysis

Gene Ontology analysis was done with the ShinyGO 0.76.2 software. Briefly, the GO Enrichment Analysis function, given a vector of genes, will return the enrichment GO categories after False Discovery Rate (FDR) control. In this analysis, we used an FDR of 0.01.

### Target-miRNA interaction analysis

For better visualization, a miRNA–mRNA network of the DE-miRNAs and relevant targets was constructed using the Cytoscape software.

### Statistical analysis

All results are expressed as mean ± SEM. One-way ANOVAs followed by Dunnett’s multiple comparisons tests or two-tailed Student′s t-test were used to detect significant differences (*p*<0.05) among treatments as indicated.

## RESULTS

### Comparison of E2Fs mRNAs and protein expression levels between hPSCs and HF arrested in G1 phase

We first sought to determine the relative abundance of canonical *E2Fs* mRNAs and proteins in hESCs, hiPSCs and, HF. Considering that the expression levels of some E2Fs fluctuate along the cell cycle and the existing differences between the percentages of hESCs, hiPSCs, and HF residing in a specific phase at any given point of the cell cycle, we opted to synchronize the studied cell types in early G1. To accomplish this, we arrested H9 hESCs, FN2.1 hiPSCs (both grown on Geltrex-coated dishes with mTeSR1 media) and HF in early G1 by using the selective CDK4/6 inhibitor PD0332991 (PD). First, we determined the cell cycle profiles by flow cytometer analysis, measuring DNA synthesis in the S phase (BrdU labelling) and DNA content (7-AAD labelling), and observed that asynchronously growing hESCs and hiPSCs exhibited a higher percentage of cells in the S phase (52.6 ± 1.3% and 42.1 ± 2.1, respectively) (Figure 1A) than HF (12.1 ± 2.6%) (data previously shown [8]) Then, we found that PD treatment led to a marked increase in the number of cells transiting the G1 phase and a decrease in the G2 population in both hPSC lines (Figure 1A) and in HF (data previously shown [8]). Next, we analyzed the mRNAs expression levels of the canonical E2F activators *E2F1*, *E2F2,* and *E2F3A* and E2F repressors *E2F3B*, *E2F4,* and *E2F5* by RT-qPCR in PD-treated cell types (hESCs, hiPSCs and HF). Importantly, we found that most *E2F* mRNA expression levels were higher in hPSCs arrested in G1 than in HF (reference of a somatic cell type). Moreover, we also quantified the mRNA expression levels of the canonical E2Fs relative to E2F1 within each cell line. In this analysis, we observed that *E2F4* had the highest expression levels, while *E2F2* had lower expression levels than other E2F activators (*E2F1* and *E2F3A*) in HF. In contrast, in the case of FN2.1 hiPSCs and H9 hESCs, E2F repressors (*E2F3B*, *E2F4* and *E2F5*) were higher expressed than most E2F activators, except for *E2F2* (its expression levels were comparable to the ones of the E2F repressors and higher than *E2F1* and *E2F3A*) (Figure 1B and Supplementary Figure S1). Finally, we measured and compared the protein expression levels of the same canonical E2Fs by western blot in early G1-arrested hESCs, hiPSCs and HF cell types. We found a good correlation between *E2F2*, *E2F3a,* and *E2F5* transcript and protein levels. On the other hand, *E2F1* and *E2F4* protein expression levels were higher in PD-arrested HF compared to hPSCs (Figure 1C).

**Figure 1.**
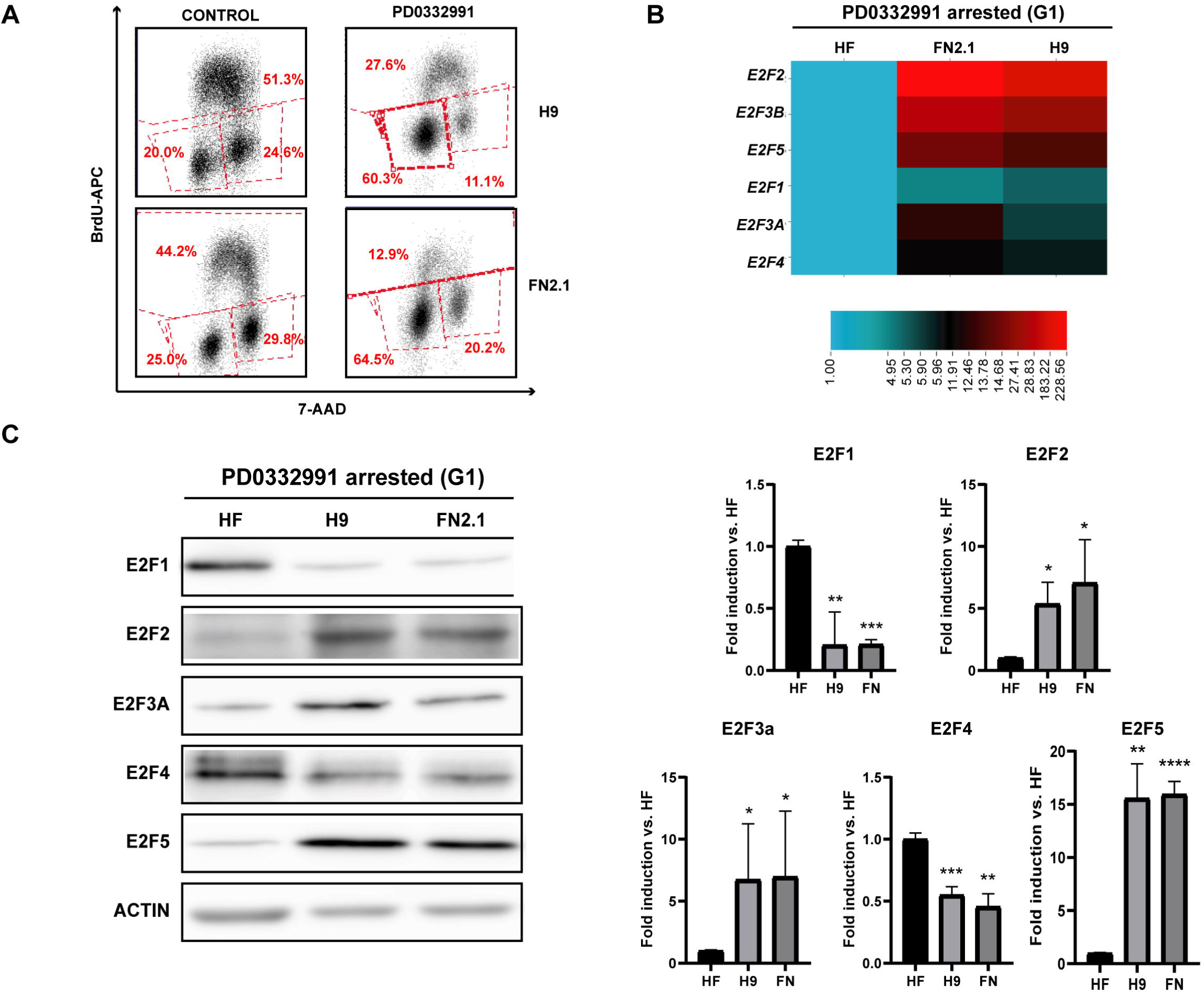
Analysis of E2F1, E2F2, E2F3A, E2F3B, E2F4 and E2F5 mRNA and protein expression levels in G1-synchronized hPSCs and HF populations. **(A)** H9 hESCs and FN2.1 hiPSCs were arrested in G1 with PD0332991 (PD arrested, 30h 5⎧M). The cell cycle profile of asynchronous (CONTROL) and pharmacologically arrested cells was analyzed after cells were stained with a 30-minute BrdU pulse and 7-AAD solution. The cell cycle positions and DNA synthetic activities of cells were determined by analyzing the correlated expression of total DNA content measured with 7-AAD and incorporated BrdU levels. Fluorescence intensity was determined with a flow cytometer. A representative BrdU plot (cell-incorporated BrdU with anti-BrdU APC vs. total DNA content with 7-AAD) is shown for each condition (n = 3). The percentage of cells in each cell cycle phase was calculated by the Accuri C6 software. **(B)** Analysis of mRNA expression levels by real-time RT-PCR in H9 hESCs, FN2.1 hiPSCs and HF following treatment with PD with primers that amplified *E2F1*, *E2F2*, *E2F3A*, *E2F3B*, *E2F4* and *E2F5* mRNAs. *RPL7* expression was used as a normalizer. The mRNA fold induction is relative to HF control cells (synchronous cells) arbitrarily set as 1. Results are shown as a *heatmap* generated with the software CIMminer. **(C)** Protein expression levels of E2F transcription factors (*E2F1, E2F2, E2F3A, E2F4* and *E2F5*) were analyzed by western blot in HF, H9 (hESCs) and FN2.1 (hiPSCs) arrested in G1 with PD0332991 (PD) (1 µΜ for 48 h for HF and 5 μM for 30h for hPSCs). Actin was used as a loading control. Mean + SEM fold induction relative to HF was graphed and representative blots of at least three independent experiments are shown. Statistical analysis was done by Student’s t-test, (*) *p* < 0.05, (**) *p* < 0.01 and (***) *p* < 0.001 vs. HF.

### Inhibition of E2Fs by pan-E2F inhibitor HLM006474

We next studied the effects of E2F inhibition on hPSCs. We use the pan-E2F inhibitor HLM006474 (iE2F). Particularly, this compound inhibits the DNA binding of all E2F complexes and consequently blocks the transcriptional activity of E2Fs. Importantly, we have previously demonstrated the efficacy of this inhibitor in hPSCs as we found that *CYCLIN E1* mRNA, a well-known target of E2Fs, was downregulated upon HLM006474 treatment [8].

First, changes in the percentage of hPSCs residing in the different cell cycle phases were studied by propidium iodide (PI) staining followed by flow cytometry analysis. Interestingly, we observed that 24 hours of 20 μM and 40 μM HLM006474 treatments increased the number of H9 hESCs in the G1 phase. Moreover, the same trend of a decrease, although not significant, was observed in the number of cells in the S phase in these cells with 24 hours of 20 μM and 40 μM iE2F treatment. On the other hand, these treatments significantly decreased the number of FN2.1 hiPSCs in the S phase, and there was a trend of an increase in the percentage of cells in the G1 phase as in hESCs (Figure 2A).

**Figure 2.**
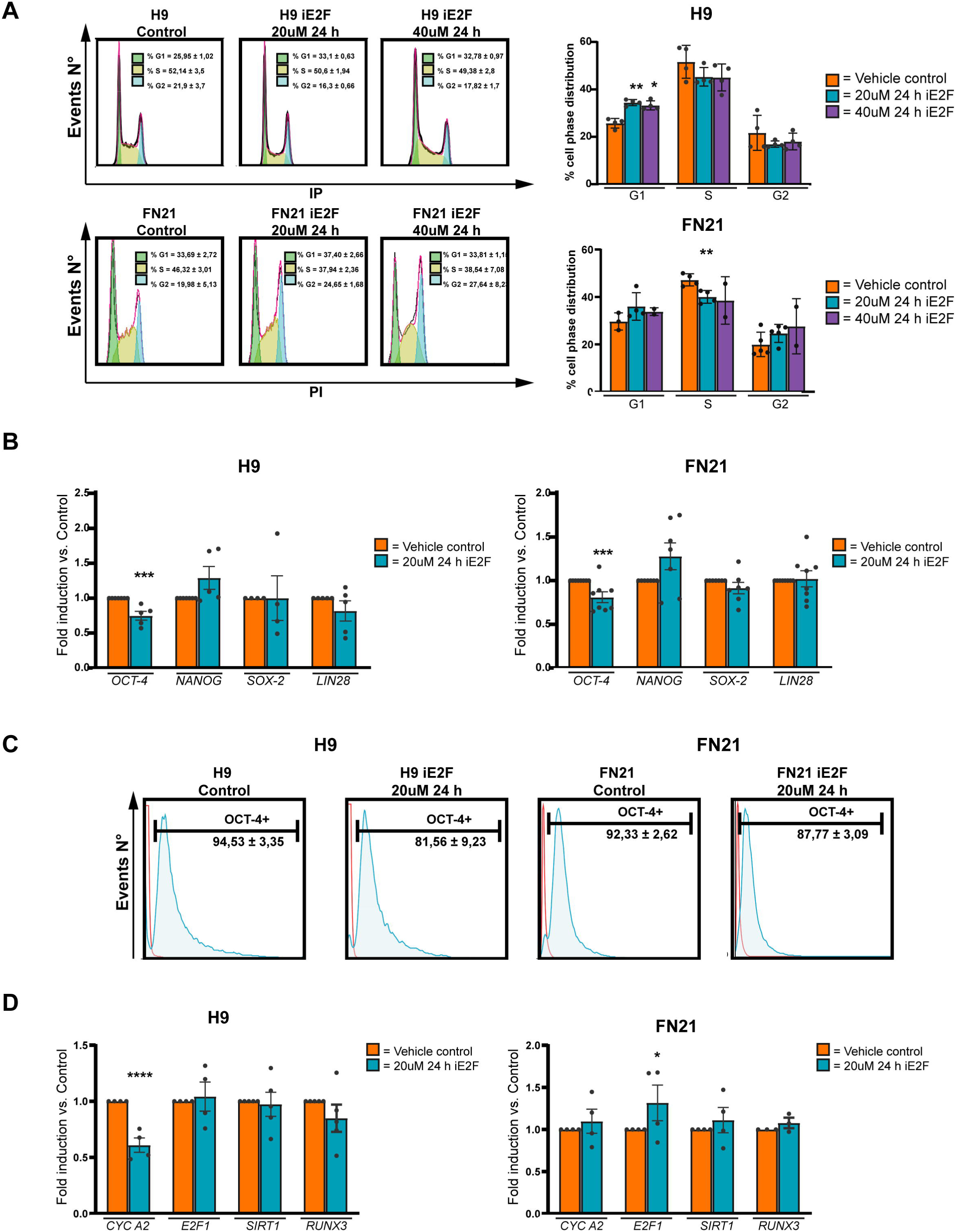
Effect of E2F inhibition on cell cycle profile and gene expression in hPSCs. **(A)** Representative histograms of three independent experiments of Propidium iodide (PI) stained H9 and FN2.1 unfixed cells treated for 24 h with HLM006474 (iE2Fs) (20 and 40 μM). The percentage of cells in each cell cycle phase was calculated by FlowJo v10.0.7’s univariate platform. Bar graphs show the percentage of cells in each cell cycle phase after 24 h of HLM006474 (iE2Fs) treatment (20 and 40μM). Mean + SEM from three independent experiments are shown. Statistical analysis was done by Student’s t-test, (*) *p*<0.05 and (**) *p*<0.01 vs. hPSCs vehicle (DMSO) control (orange bar). **(B)** Analysis of *OCT-4, NANOG, SOX-2* and *LIN28* mRNA expression levels, quantified by RT-qPCR, in iE2Fs-treated H9 hESCs and FN2.1 hiPSCs. *RPL7* expression was used as a normalizer. Mean ± SEM fold induction relative to vehicle control hPSCs (orange bars) (arbitrarily set as 1) from at least four independent experiments is graphed. Vehicle: DMSO. Statistical analysis was done by Student’s t-test, (***) p<0.001 vs vehicle control hPSCs. **(C)** Flow cytometric analysis of control and iE2Fs treated hPSCs (24h 20 μM) showing the percentage of OCT-4^+^ cells. Representative histograms are shown. The red tracing shows the secondary antibody-only control. **(D)** Analysis of *CYCLIN A2, E2F1, SIRT1* and *RUNX3* mRNA expression levels, quantified by RT-qPCR in iE2Fs-treated H9 hESCs and FN2.1 hiPSCs (24h, 20 μM)*. RPL7* expression was used as a normalizer. Mean ± SEM fold induction relative to vehicle control hPSCs (orange bars) (arbitrarily set as 1) from at least three independent experiments is graphed. Vehicle: DMSO. Statistical analysis was done by Student’s t-test, (*) p<0.05; p<0.0001 (****) vs vehicle control hPSCs.

Then, we quantified the percentage of cell viability after 24 and 48 hours of HLM006474 incubation (20 and 40 μM) using an XTT/PMS vital dye assay. We found that under the mentioned experimental conditions cell viability slightly decreased in H9 hESCs with 40 μM at 24 and 48 hours of treatment (Supplementary Figure S2A). Similar results were observed when living or dead cells were counted using Trypan blue dye at 48 hours of treatment with a 40 μM concentration (Supplementary Figure S2B). On the other hand, we observed a slight decrease in FN2.1 hiPSC viability under 20 and 40 μM HLM006474 at 24 and 48 hours of treatment with an XTT/PMS vital dye assay. Surprisingly, no difference in cell death was found between the different treatments when we used the Trypan blue dye in FN2.1 hiPSCs (Supplementary Figure S2B). Finally, we measured cell death by flow cytometry analysis of PI staining. Importantly, only cells exhibiting loss of plasma membrane integrity (late apoptosis or necrosis) are stained with PI. Strikingly, no difference in cell death was found between the different treatments when we used the PI dye in either hESCs or hiPSCs (Supplementary Figure S2C).

Thus, we chose 20 μM for 24 hours of HLM006474 treatment for further experiments as it was the lowest concentration tested and temporal period that induced changes in the cell cycle profile without or only slightly affecting hPSC viability. Interestingly, we determined a downregulation, quantified by RT-qPCR, of the mRNA expression levels of the stemness marker *OCT-4,* in H9 hESCs and FN2.1 hiPSCs treated with 20 μM HLM006474 for 24 hours. On the contrary, no significant changes in the mRNA expression levels of other master transcription factors that control pluripotency, such as *NANOG*, *SOX-2* and *LIN28,* were observed in iE2F-treated hPSCs (Figure 2B). Besides, concomitantly to the downregulation of the mRNA levels, a reduction in OCT-4 fluorescent signal levels was detected by flow cytometry analysis in H9 hESCs and FN2.1 hiPSCs treated with iE2F (20 μM, 24 h) (Figure 2C).

Furthermore, to evaluate the effect of HLM006474 treatment (20 μM; 24 hours) in hPSCs we selected a group of genes that were previously reported to be regulated by E2F transcription factors in other biological models, such as *CYCLIN A2*, *E2F1*, *SIRT1* and *RUNX3*, and analyzed their mRNA expression levels by RT-qPCR in. Results shown in Figure 2D indicate that *CYCLIN A2* mRNA expression levels decreased in iE2F-treated H9 hESCs and that E2F1 mRNA expression levels increased in iE2F-treated FN21 hiPSCs.

### High throughput small RNA-sequencing and differential expression analysis

Next, we aimed to identify E2Fs-regulated miRNAs in hPSCs. In this sense, we performed and compared massive sequencing analysis of small RNAs of H9 hESCs untreated (control) and treated with HLM006474 20 μM for 24 hours. From our sequencing results, we first analyzed the miRNA levels profile of all replicates of H9 hESCs either untreated (control) or treated with the E2Fs pan-inhibitor. We were able to detect 885 microRNAs using 10 reads in one of the samples as a cut-off point. The *heatmap* shows a similar miRNA expression distribution between replicates and both types of samples (Supplementary Figure S3A).

Then, we performed a differential expression analysis to elucidate whether HLM006474 treatment induces changes in the expression levels of miRNAs. First, we performed some diagnostic plots to corroborate that the analysis was correct. We determined that the normalization made by the used packages (EGER and limma) was adequate (Supplementary Figure S3B). Principal component analysis (PCA) supported what was seen in the *heatmap* as it shows a small distance between samples from both populations (57% of variance explained) (Supplementary Figure S3C). Finally, we did a Mean-Average plot (MA-plot). The most interesting miRNA candidates are observed in the plot’s upper/lower right quadrant (Supplementary Figure S3D). From a differential expression analysis, we found that 52 miRNAs were differentially expressed with a 0.1 FDR and 1.5-fold change between H9 hESCs untreated and treated with the E2Fs inhibitor (Volcano plot, Figure 3B and Supplementary Table S4). Twenty miRNAs were selected from the original list of 52 as interesting candidates, some of them already known to be regulated by the E2F transcription factors family and others with a good amount of reads whose relationship with these factors or with the cell cycle in hESCs has not yet been reported. The table shows Fold change and *p*-values of the 10 most up (green) and down-regulated (red) miRNAs in H9 hESCs iE2F treated cells (Figure 3A).

**Figure 3.**
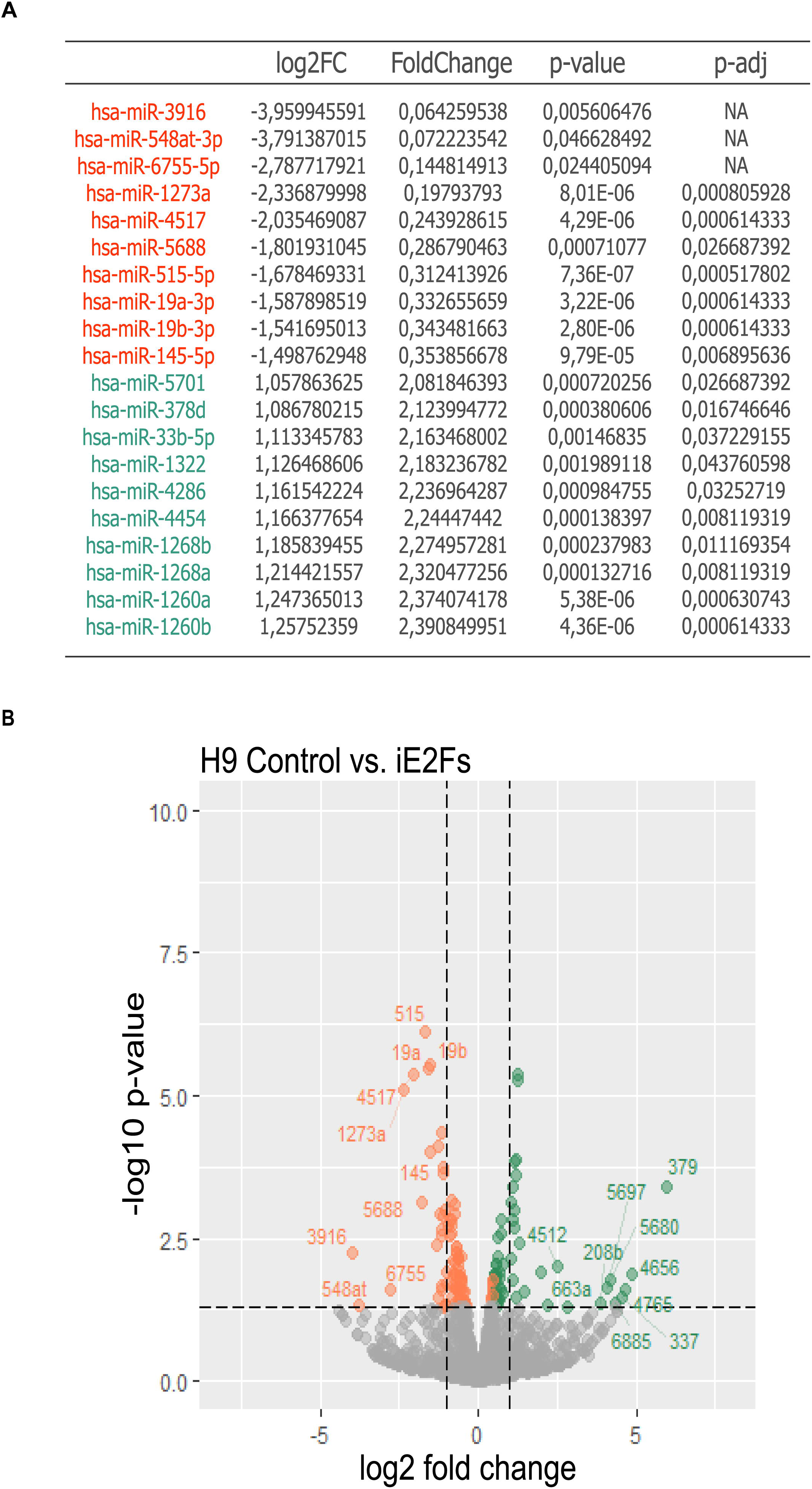
Differential expression analysis. A differential expression analysis was performed between control and E2Fs pan-inhibitor HLM006474-treated (24 h, 20 μM; iE2Fs) H9 hESCs using the DESeq2 package in R with a false discovery rate (FDR) of 0.1. The design formula was set according to the treatment to analyze the effect of E2F inhibition on miRNA expression. **(A)** The table summarizes the 10 most up- (green) and down-regulated (red) miRNAs in H9 hESCs after 24 h of 20 μM pan-E2F inhibitor treatment. **(B)** Volcano-plot of changes on miRNAs expression between treatment (control vs iE2F) comparisons according to significance. The Black dotted line belongs to a 1/-1 log2 fold change and a –log10 *p*-value of 1.3/-1.3 that belongs to a 0.1 *p*-value. Down-regulated miRNAs in H9 iE2Fs are in red and Up-regulated in green. Only the miRNA numbers of the 10 most up- and down-regulated miRNAs are shown.

Later, we validated the expression levels of the selected miRNA candidates by RT-qPCR with specific stem-loop primers to confirm our RNA-seq results (Figure 4). We specifically amplify 7 miRNAs (miR-19a and b, miR-1260a and b, miR-4454, miR-454, and miR-301a) of the 20 candidates in both hPSC cell lines. From this list, we observed that miR-454-3p, miR-19a-3p, miR-19b-3p and miR-301a-3p expression levels were down-regulated in iE2Fs-treated hPSCs. On the other hand, we observed a tendency of increase in miR-4454, miR-1260a and miR-1260b expression levels (Figure 4). These changes in the expression levels found in iE2Fs-treated hPSCs relative to their untreated counterparts were like the observed from the analysis of the sequencing results.

**Figure 4.**
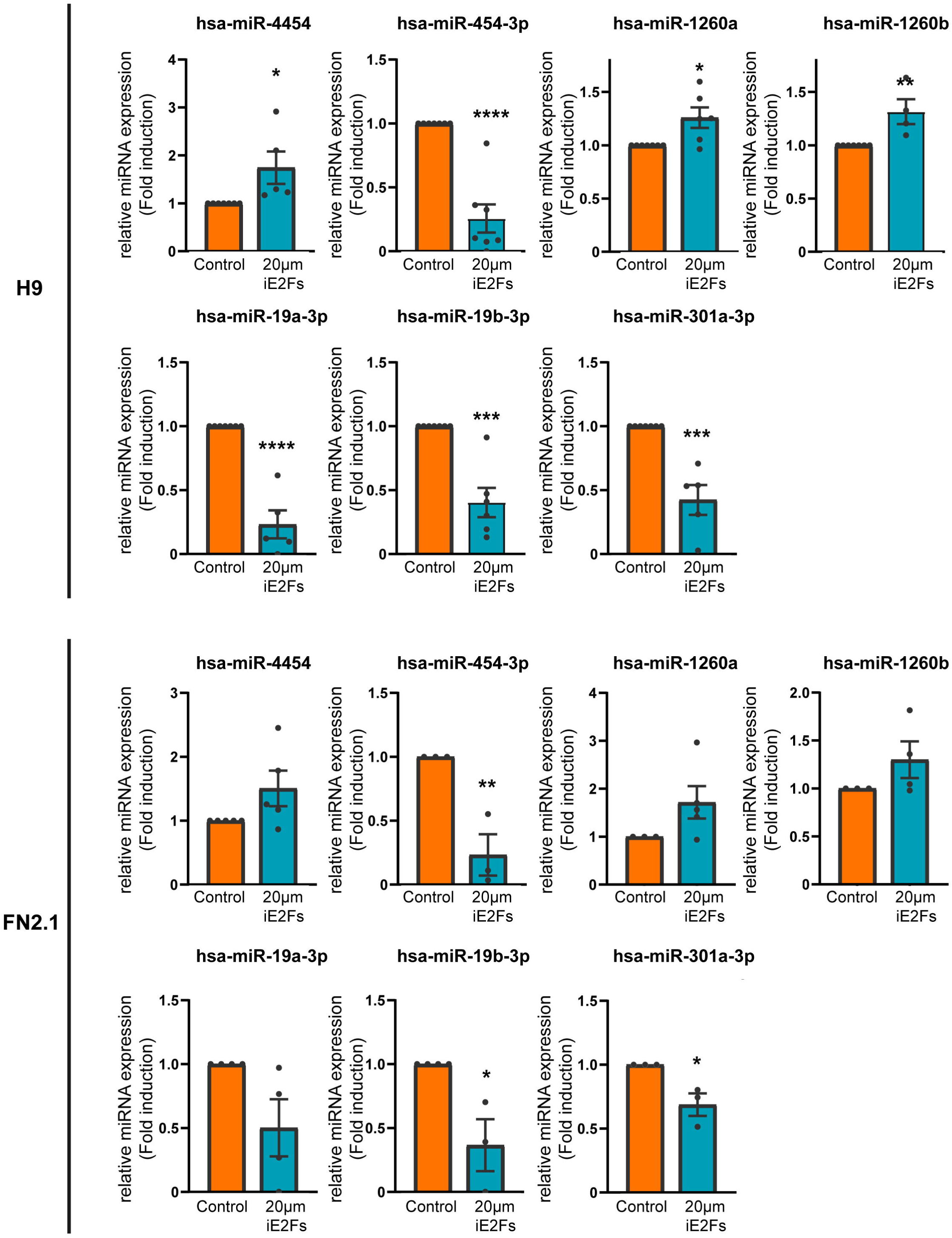
miRNAs validation. Seven miRNAs were selected after specificity primer validation from 20 differentially expressed miRNA candidates. Analysis of miR-4454, miR-1260a, miR-1260b, miR-301a-3p, miR-454-3p, miR-19a-3p, miR-19b-3p miRNA expression levels quantified by RT-qPCR with specific stem-loop primers in control and HLM006474 (24 h, 20 μM; iE2Fs) treated hPSCs. *RNU6B* expression was used as a normalizer. Graphs show mean ± SEM miRNA fold induction relative to vehicle control (arbitrarily set as 1) of at least three independent experiments. Statistical analysis was done by Student’s t-test, (*) *p*<0.05; (**) *p*<0.01; (***) *p*<0.001; (****) *p*<0.0001. iE2Fs-treated vs vehicle (DMSO) control.

### Target genes of the selected differentially expressed miRNAs

Next, we sought to explore which biological processes are potentially regulated by the 52 differentially expressed miRNAs. To this end, we analyzed the gene ontology terms (GO) associated with a list of predicted targets (10 gene targets for each miRNA with the higher *p*-binding) of 51 of these miRNAs from the miRWalk 3.0 software (no ID match was found for has-miR-1273a) (Supplementary Table S4). Importantly, the analysis of the GO terms revealed enrichment in processes related to cell cycle regulation, development, and neural differentiation (Figure 5A). This reflects that miRNAs are involved in diverse regulatory pathways. In this context, it is highly important to identify the targets of specific miRNAs to understand the mechanism of their regulation. In this sense, from the predicted genes targeted by the differentially expressed miRNAs, we plot the target-miRNA interaction network. Figure 5B shows the miRNA-gene network of 427 putative target genes. Among them, *ANKEF1*, *CDC27*, *DCUN1D3*, *DENND4C*, *KATNAL1*, *PRKAR1A* and *PTBP2* are targeted by three miRNAs. In all cases, this central regulatory molecule belongs to the miR-548 family. This is consistent with the knowledge that miRNA members of the same family share an identical 5′ terminal seed sequence and regulate the same pool of target genes. Besides, this reflects that miRNAs can regulate many target genes and target genes can be regulated by many miRNAs and, all together, how complex these regulation networks are.

**Figure 5.**
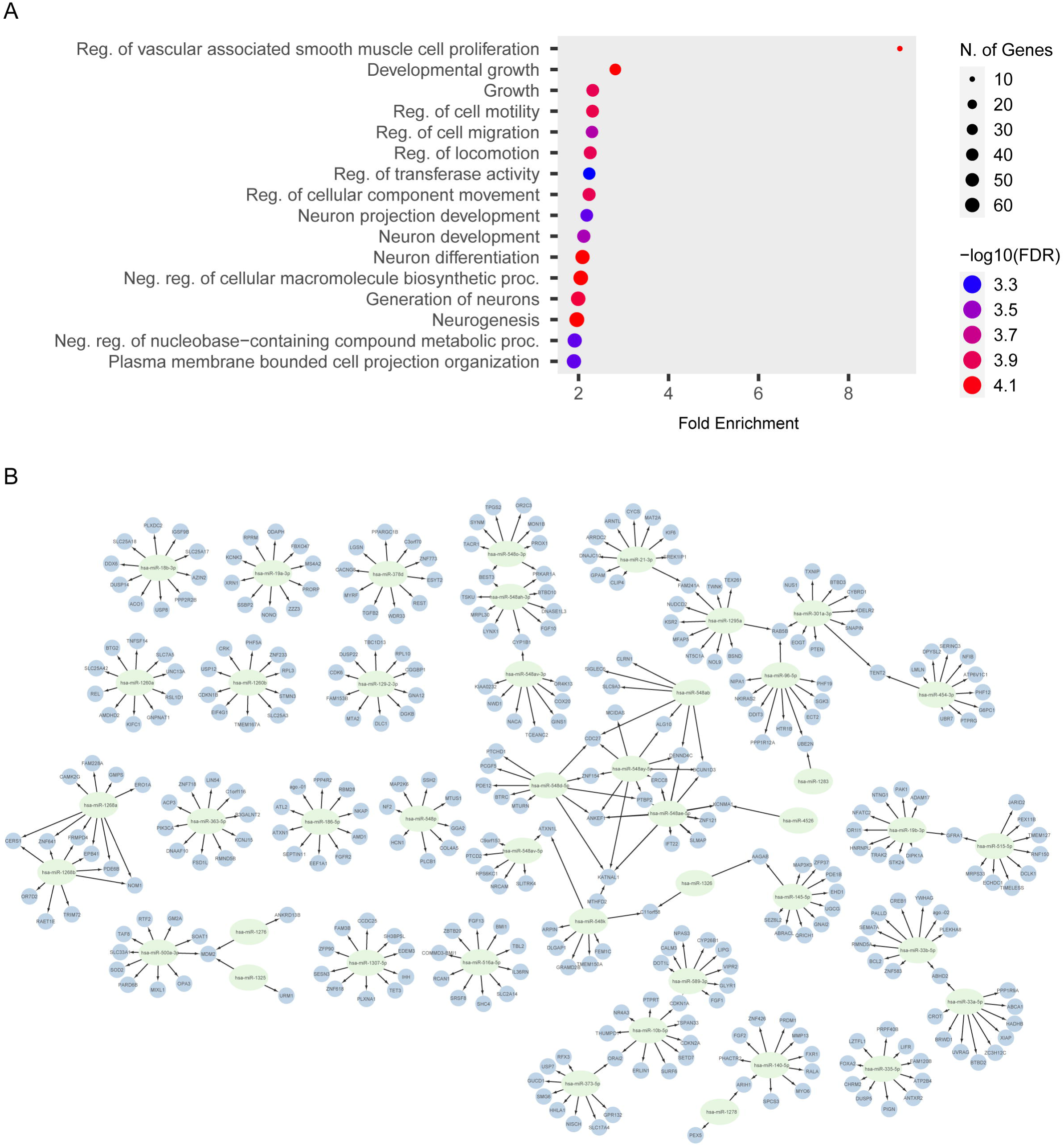
Gene Ontology analysis and interaction map. **(A)** Gene Ontology (GO) terms related to the target genes list predicted for 51 miRNAs from the differential expression analysis are shown in the dot plot (no ID match was found for has-miR-1273a). The terms are based on the biological processes level. The dot size represents the number of genes included in each GO term and the colourimetric scale is based on the p-value. Gene Ratio represents the number of genes per the total number of genes in each GO term. GO enrichment analysis was made with the ShinyGo software with a 0.05 FDR. **(B)** The interaction network shows nodes and connections between miRNA candidates and the predicted target genes. The green nodes represent the miRNA, and the light blue nodes represent its targeted gene. The opacity of the black edge links signifies the target score between the miRNA and the target gene. The network was made with the Cytoscape software.

## DISCUSSION

Cell division and cycle progression are controlled by mechanisms that ensure the proper transmission of genetic information from one generation to the next. Thus, cell cycle control is a complex process involving different signalling pathways. For years, cell cycle regulation has been studied primarily due to its relevance in hyperproliferative diseases such as cancer. Currently, interest in cell cycle regulation has expanded into other emerging areas of research, such as stem cell biology [25]. The study of hPSC properties is strongly associated with future hopes for their possible use in regenerative cell-based therapies. Because of this, a complete understanding of the molecular basis of the mechanisms involved in cell cycle progression would facilitate efforts to develop new *in vitro* propagation methods [4].

E2F transcription factors are critical for the temporal expression of cell cycle oscillating genes and non-coding RNAs. The pRB/E2F pathway is crucial for the cell cycle by stimulating G1 phase cyclins and CDKs to form complexes. In hPSCs, the roles of the different pRB/E2F complexes and the factors that determine what each complex binds to remain uncertain. There are few reports on the expression of E2F transcription factors in pluripotent cells. Herein, we determined that G1-arrested hPSCs express higher mRNA levels of canonical *E2F1*, *E2F2, E2F3A* and *B, E2F4* and *E2F5* mRNAs than somatic cells like HF. Besides, we also found it particularly interesting that, when we analyzed the E2F mRNA expression levels within each cell line, we observed that E2F repressors (E2F3B, E2F4, and E2F5) were higher expressed than E2F activators (except for E2F2) in both human pluripotent cell lines (H9 hESCs and FN2.1 hiPSCs). This is consistent with a previous report by Becker *et al.* in hESCs where they demonstrated that repressor E2Fs are predominantly expressed in these cells as opposed to somatic cells, being E2F5 the most abundant member of its class in hESCs. Moreover, they found that E2F1, E2F2, and E2F3 together account for only 17–24% of total E2F activity in H1 and H9 hESCs, but as much as 58% in human IMR90 fibroblasts [25].

At a protein level, although most canonical E2F exhibited higher expression levels in G1-arrested hPSCs relative to HF, we observed the opposite in the case of E2F1 and E2F4. Interestingly, Hsu *et al*. recently reported that in mouse embryonic stem cells (mESCs) E2F4 acts as a transcriptional activator that promotes the expression of cell cycle genes independently of the pRB family instead of as a transcriptional repressor. Hence, the activity of E2F4 is critical for mESCs proliferation and survival [26]. These results reflect versatility in the roles achieved by cell cycle regulators that may rely on the molecular settings displayed by somatic and pluripotent stem cells. Furthermore, it is not yet known whether E2F4 functions as a transcription activator or repressor in hPSCs and may explain the inconsistencies between mRNA and protein expression levels for this E2F.

Previous reports pointed to a link between the cell cycle and mammalian stem cell pluripotency [27]. A study demonstrates that following ablation of G1 cyclins, mESCs attenuated their pluripotent characteristics. Particularly, this resulted in a down-regulation of OCT-4, NANOG and SOX-2 at the protein level. They established that G1 cyclins, including CYCLIN E1, together with their associated CDKs, phosphorylate and stabilize the core pluripotency factors NANOG, SOX-2 and OCT-4 [27]. Also, in line with this, Li *et al.* demonstrated that E2F inhibition with HLM006474 (30 μM, 24 hours) increased hESCs (line HUES6) differentiation across all germ layers promoting enhanced expression of several lineage-specific markers [28]. In this sense, we observed in hESCs and hiPSCs that E2F inhibition with HLM006474 resulted in a decrease in *OCT-4* mRNA expression levels and the percentage of OCT-4^+^ hPSCs. Interestingly, *CYCLIN E1* is transcriptionally activated by E2F transcription factors and we had previously shown that pan-E2F treatment leads to a down-regulation of *CYCLIN E1* mRNA expression levels in hPSCs [8]. Further experiments should be performed to elucidate if the changes that we observed in OCT-4 expression levels are linked to the down-regulation of CYCLIN E1 induced by E2F inhibition.

On the other hand, miRNAs fine-tune numerous functions in pluripotent stem cells, such as pluripotency, self-renewal and differentiation [29]. These processes must be spatially and temporally regulated so that pluripotent cells can correctly develop and warrant accurate embryogenesis during development. As previously mentioned, miRNAs are relevant gene regulatory factors that bind to target genes and inhibit their expression at the post-transcriptional level, thus managing protein output such as those that regulate the cell division cycle. To date, how miRNAs are regulated in hPSCs remains poorly understood. Several clusters and single miRNAs that control self-renewal and pluripotency have been identified. These include miR-290-295, miR-302, miR-17-92, miR-106b-25 and miR-106a-363, which are functionally up-regulated to suppress negative regulators and enhance pluripotent transcription factors such as NANOG and MYC [30, 31].

E2F transcription factors had been identified as both targets of miRNAs and regulators of their expression. In this regard, a number of E2F-regulated miRNAs have been determined, including the miR-17-92, miR-106a-363, and miR-106b-25 clusters as well as miR-449a and miR-449b. Remarkably, some of these E2F-regulated miRNAs modulate E2Fs and/or other components of the pRB/E2F pathway, establishing a regulatory loop that controls E2F activity [32]. Besides, by ChIP assays using a pool of fibroblast-derived from mouse embryos Bueno *et al.* have identified direct binding of E2Fs transcription factors to the regulatory sequences of four genes, expressing the let-7a-7d, let-7i, miR-15b-16-2, or miR-106b-25 transcripts, during the G1/S transition [33]. These G1-related miRNAs are induced by E2F1 or E2F3A and are down-regulated in E2F1-knockout or E2F3-knockdown cells. These results suggest the functional relevance of these E2F transcription factors in the control of these miRNAs [28, 33].

In this sense, to determine miRNAs regulated by E2F transcription factors in hPSCs, we treated cells with the E2F inhibitor (pan-E2F inhibitor) HLM006474. Exposure to this inhibitor led to an increase in the percentage of cells residing in the G1 phase in both hPSCs lines without affecting (at certain concentrations) cell viability. It has been reported that each member of the E2F family have unique and specialized functions and, besides, a down-regulation of total intracellular E2F activity can cause apoptosis, growth arrest, or both [34]. Moreover, several studies suggest that E2Fs must bind to DNA to stimulate proliferation, which is supported by our observations since HLM006474 inhibits DNA binding of E2F complexes [35, 36]. Herein, we determined that miR-19a-3p, miR-19b-3p, miR-4454, miR-1260a, miR-1260b, miR-454-3p and miR-301a-3p are regulated by E2Fs. Intriguingly, using TransmirR v20, an updated transcription factor-miRNA regulation database, we found that some of the selected miRNAs own putative binding motifs of E2Fs within their promoter region.

Lu *et al* observed that repression of miR-301a-3p (one of the differentially expressed miRNAs) suppressed cancer cell growth, proliferation and arrested cell cycle at the G0/G1 phase in Laryngeal Squamous Cell Carcinoma, suggesting that this miRNA may be regulated by transcription factors involved in cell cycle regulation [37]. Moreover, supporting this idea miR-301a-3p was also reported to regulate proliferation through its target gene Phosphatase and tensin homolog (PTEN) in esophageal squamous cells [38], and Run-related transcription factor 3 (RUNX3) in prostate cancer [39]. Interestingly, we observed a decrease in the expression levels of this miRNA in iE2F-treated hPSCs where, in turn, we saw an increase in the percentage of cells residing in the G1 phase.

From the remaining differentially expressed miRNAs, we highlight miR-19a and miR-19b. It has been described that E2F1, E2F2 and E2F3 bind directly to the promoter of the *C13orf25* gene (where the cluster that encodes miR-19a and miR-19b-1 is located) activating its transcription in HeLa cells. Also, E2F1 can activate the transcription of the miR-106a-303 cluster (from where miR-19-b-2 is encoded) [40]. Even though miR-19b-1 and miR-19b-2 are transcribed from different genes, their mature sequence is the same and it is commonly named as miR-19b. This information is in line with our results where we observed in HLM006474 treated cells, a decrease in the expression of this miRNA. This suggests that these miRNAs would be regulated by some of the E2F transcription factor(s) in hPSCs.

As for the other differentially expressed miRNAs, we found no reports on their relationship with E2F transcription factors or pluripotent stem cells. In particular, it was described that miR-1260b has Secreted Frizzled-related protein 1 *(SFRP1*) as a target gene, and through its silencing, activates the Wnt/β-catenin pathway in an adenocarcinoma model [41]. It has also been described that miR-454-3p targets this gene and is capable of activating this pathway and promoting metastasis in breast cancer [42]. It would be interesting to deepen their study and determine which pathways these and the above-mentioned miRNAs regulate in our experimental model. However, this aim may be challenging given that the changes in the expression of these miRNAs because of the pan-iE2F were subtle and compensation effects may occur among them.

Thus, the observed biological effect may be a consequence of the activities of multiple interrelated miRNAs. Therefore, given that a miRNA may regulate more than one target gene and different miRNAs may also regulate the same target genes [43], different combinations of inhibition or overexpression of these miRNAs should be made to find a relevant biological effect.

Increasing evidence suggests that miRNAs are pivotal regulators of development and cellular homeostasis through their control of diverse biological processes [43]. In this sense, gene ontology (GO) analysis of the predicted target genes of the DE miRNAs suggests that they would be regulating genes, directly or indirectly, involved in the cell cycle and different pathways of differentiation. The GO terms also include neuronal differentiation and development. Several studies have shown that members of the miR-19 family, among which miR-19a and miR-19b stand out, play an important role in the regulation of the development of the nervous system, cardiovascular system, and blood vessel formation, among others. In addition, a key role in neuronal differentiation, survival and migration has been described [40]. It is well known that cell cycle regulation and differentiation processes are intrinsically related. These results suggest that these miRNAs, which could be regulated directly or indirectly by the E2F transcription factors, may participate in this balance thus affecting the observed G1 phase alteration in human pluripotent stem cells.

## ACKNOWLEDGMENTS

This work was supported by grants from ANPCyT PICT-2016-2621 and Fundación FLENI. The authors would like to thank Marcela Cañari for her skilful assistance.

## REFERENCES

1. Takahashi, K., et al., Induction of pluripotent stem cells from adult human fibroblasts by defined factors. Cell, 2007. 131(5): p. 861–72.

2. Thomson, J.A., et al., Embryonic stem cell lines derived from human blastocysts. Science, 1998. 282(5391): p. 1145–7.

3. Klimanskaya, I., N. Rosenthal, and R. Lanza, Derive and conquer: sourcing and differentiating stem cells for therapeutic applications. Nat Rev Drug Discov, 2008. 7(2): p. 131–42.

4. Abdelalim, E.M., Molecular mechanisms controlling the cell cycle in embryonic stem cells. Stem Cell Rev Rep, 2013. 9(6): p. 764–73.

5. White, J. and S. Dalton, Cell cycle control of embryonic stem cells. Stem Cell Rev, 2005. 1(2): p. 131–8.

6. Becker, K.A., et al., Self-renewal of human embryonic stem cells is supported by a shortened G1 cell cycle phase. J Cell Physiol, 2006. 209(3): p. 883–93.

7. Ghule, P.N., et al., Reprogramming the pluripotent cell cycle: restoration of an abbreviated G1 phase in human induced pluripotent stem (iPS) cells. J Cell Physiol, 2011. 226(5): p. 1149–56.

8. Rodriguez Varela, M.S., et al., Regulation of cyclin E1 expression in human pluripotent stem cells and derived neural progeny. Cell Cycle, 2018. 17(14): p. 1721–1744.

9. Filipczyk, A.A., et al., Differentiation is coupled to changes in the cell cycle regulatory apparatus of human embryonic stem cells. Stem Cell Res, 2007. 1(1): p. 45–60.

10. Sela, Y., et al., Human embryonic stem cells exhibit increased propensity to differentiate during the G1 phase prior to phosphorylation of retinoblastoma protein. Stem Cells, 2012. 30(6): p. 1097–108.

11. Thurlings, I. and A. de Bruin, E2F Transcription Factors Control the Roller Coaster Ride of Cell Cycle Gene Expression. Methods Mol Biol, 2016. 1342: p. 71–88.

12. Carleton, M., M.A. Cleary, and P.S. Linsley, MicroRNAs and cell cycle regulation. Cell Cycle, 2007. 6(17): p. 2127–32.

13. Wang, Y., et al., Embryonic stem cell-specific microRNAs regulate the G1-S transition and promote rapid proliferation. Nat Genet, 2008. 40(12): p. 1478–83.

14. Rodriguez-Varela, M.S., et al., miR-302 family, miR-145 and miR-296 temporal expression profile along the cell cycle of human pluripotent stem cells. Gene Expr Patterns, 2021. 40: p. 119168.

15. Julian, L.M. and A. Blais, Transcriptional control of stem cell fate by E2Fs and pocket proteins. Front Genet, 2015. 6: p. 161.

16. Beijersbergen, R.L., et al., Regulation of the retinoblastoma protein-related p107 by G1 cyclin complexes. Genes Dev, 1995. 9(11): p. 1340–53.

17. Lv, Y., et al., E2F8 is a Potential Therapeutic Target for Hepatocellular Carcinoma. J Cancer, 2017. 8(7): p. 1205–1213.

18. Neganova, I. and M. Lako, G1 to S phase cell cycle transition in somatic and embryonic stem cells. J Anat, 2008. 213(1): p. 30–44.

19. Becker, K.A., et al., Human embryonic stem cells are pre-mitotically committed to self-renewal and acquire a lengthened G1 phase upon lineage programming. J Cell Physiol, 2010. 222(1): p. 103–10.

20. Questa, M., et al., Generation of iPSC line iPSC-FH2.1 in hypoxic conditions from human foreskin fibroblasts. Stem Cell Res, 2016. 16(2): p. 300–3.

21. Romorini, L., et al., Effect of antibiotics against Mycoplasma sp. on human embryonic stem cells undifferentiated status, pluripotency, cell viability and growth. PLoS One, 2013. 8(7): p. e70267.

22. Mucci, S., et al., Acute severe hypoxia induces apoptosis of human pluripotent stem cells by a HIF-1alpha and P53 independent mechanism. Sci Rep, 2022. 12(1): p. 18803.

23. Isaja, L., et al., Chemical hypoxia induces apoptosis of human pluripotent stem cells by a NOXA-mediated HIF-1alpha and HIF-2alpha independent mechanism. Sci Rep, 2020. 10(1): p. 20653.

24. Romorini, L., et al., AKT/GSK3beta signaling pathway is critically involved in human pluripotent stem cell survival. Sci Rep, 2016. 6: p. 35660.

25. Barta, T., et al., Cell cycle regulation in human embryonic stem cells: links to adaptation to cell culture. Exp Biol Med (Maywood), 2013. 238(3): p. 271–5.

26. Hsu, J., et al., E2F4 regulates transcriptional activation in mouse embryonic stem cells independently of the RB family. Nat Commun, 2019. 10(1): p. 2939.

27. Liu, L., et al., G1 cyclins link proliferation, pluripotency and differentiation of embryonic stem cells. Nat Cell Biol, 2017. 19(3): p. 177–188.

28. Li, J., et al., A transient DMSO treatment increases the differentiation potential of human pluripotent stem cells through the Rb family. PLoS One, 2018. 13(12): p. e0208110.

29. Divisato, G., et al., The Key Role of MicroRNAs in Self-Renewal and Differentiation of Embryonic Stem Cells. Int J Mol Sci, 2020. 21(17).

30. Li, N., et al., microRNAs: important regulators of stem cells. Stem Cell Res Ther, 2017. 8(1): p. 110.

31. Maraghechi, P., et al., Pluripotency-Associated microRNAs in Early Vertebrate Embryos and Stem Cells. Genes (Basel), 2023. 14(7).

32. Ofir, M., D. Hacohen, and D. Ginsberg, MiR-15 and miR-16 are direct transcriptional targets of E2F1 that limit E2F-induced proliferation by targeting cyclin E. Mol Cancer Res, 2011. 9(4): p. 440–7.

33. Bueno, M.J., et al., Multiple E2F-induced microRNAs prevent replicative stress in response to mitogenic signaling. Mol Cell Biol, 2010. 30(12): p. 2983–95.

34. Ma, Y., et al., A small-molecule E2F inhibitor blocks growth in a melanoma culture model. Cancer Res, 2008. 68(15): p. 6292–9.

35. Wu, C.L., et al., Expression of dominant-negative mutant DP-1 blocks cell cycle progression in G1. Mol Cell Biol, 1996. 16(7): p. 3698–706.

36. Maehara, K., et al., Reduction of total E2F/DP activity induces senescence-like cell cycle arrest in cancer cells lacking functional pRB and p53. J Cell Biol, 2005. 168(4): p. 553–60.

37. Lu, Y., et al., Hsa-miR-301a-3p Acts as an Oncogene in Laryngeal Squamous Cell Carcinoma via Target Regulation of Smad4. J Cancer, 2015. 6(12): p. 1260–75.

38. Zhang, N. and J.F. Liu, MicroRNA (MiR)-301a-3p regulates the proliferation of esophageal squamous cells via targeting PTEN. Bioengineered, 2020. 11(1): p. 972–983.

39. Fan, L., et al., MicroRNA⍰301a⍰3p overexpression promotes cell invasion and proliferation by targeting runt⍰related transcription factor 3 in prostate cancer. Mol Med Rep, 2019. 20(4): p. 3755–3763.

40. Li, X., et al., miR-19 family: A promising biomarker and therapeutic target in heart, vessels and neurons. Life Sci, 2019. 232: p. 116651.

41. Ren, J., et al., miR-1260b Activates Wnt Signaling by Targeting Secreted Frizzled-Related Protein 1 to Regulate Taxane Resistance in Lung Adenocarcinoma. Front Oncol, 2020. 10: p. 557327.

42. Ren, L., et al., MiR-454-3p-Mediated Wnt/beta-catenin Signaling Antagonists Suppression Promotes Breast Cancer Metastasis. Theranostics, 2019. 9(2): p. 449–465.

43. Liu, B., J. Li, and M.J. Cairns, Identifying miRNAs, targets and functions. Brief Bioinform, 2014. 15(1): p. 1–19.

