## Supplementary information for "“Identification of microRNAs regulated by E2F transcription factors in human pluripotent stem cells”"

**Author affiliations:**

**Supplementary methods:**

***Antibodies and primers used:***

|  | **Primer sequence (5'** → **3')** | |
| --- | --- | --- |
| **Name** | **Forward** | **Reverse** |
| *RPL7* | AATGGCGAGGATGGCAAG | TGACGAAGGCGAAGAAGC |
| *E2F1* | ATGTTTTCCTGTGCCCTGAG | ATCTGTGGTGAGGGATGAGG |
| *E2F2* | TAAGGAGCAGACAGTGATTG | AGAGGGTGGAGGTAGAGG |
| *E2F3A* | GCCCCGATTATTTTTGGCCC | TCATCTCTCGCTCCTGCTCT |
| *E2F3B* | TACAGCAAGCAGGCAAAGC | GTGAGCAGACCAAGAGACG |
| *E2F4* | CATAGGGGGCAGTGTCTTGT | CTAAAGGCCCAGCAGAAGTG |
| *E2F5* | GGACCTATCCATGTGCTGCT | TGTTGCTCAGGCAGATTTTG |
| *OCT-4* | CTGGGTTGATCCTCGGACCT | CACAGAACTCATACGGCGGG |
| *NANOG* | AAAGAATCTTCACCTATGCC | GAAGGAAGAGGAGAGACAGT |
| *SOX-2* | AGCATGGAGAAAACCCGGTACGC | CGTGAGTGTGGATGGGATTGGTGT |
| *LIN28* | TCAGGCTTGGGTTCACACCATCAC | GGTTGCCCCCAGAACCCTCAC |
| *CYC A2* | CCTGCAAACTGCAAAGTTGA | AAAGGCAGCTCCAGCAATAA |
| *SIRT-1* | CCTGTGAAAGTGATGAGGAGGA | GAATTGTTCGAGGATCTGTGCC |
| *RUNX3* | AGGCAATGACGAGAACTACTCC | CGAAGGTCGTTGAACCTGG |

**Supplementary Table S1.** *Primers used for RT-qPCR experiments.*

|  | **Primer sequence ( 5'** → **3')** | |
| --- | --- | --- |
| **Name** | **SLO primer** | **Forward** |
| *mir-4454* | *GTCGTATCCAGTGCAGGGTCCGAGGTATTCGCACTGGATACGACTGGTGC* | *GGATCCGAGTCACGGCACCA* |
| *mir-454-3p* | *GTCGTATCCAGTGCAGGGTCCGAGGTATTCGCACTGGATACGACACCCTA* | *TAGTGCAATATTGCTTATAGGGT* |
| *mir-1260a* | *GTCGTATCCAGTGCAGGGTCCGAGGTATTCGCACTGGATACGACTGGTGG* | *ATCCCACCTCTGCCACCA* |
| *mir-1260b* | *GTCGTATCCAGTGCAGGGTCCGAGGTATTCGCACTGGATACGACATGGTG* | *ATCCCACCACTGCCACCAT* |
| *mir-19a-3p* | *GTCGTATCCAGTGCAGGGTCCGAGGTATTCGCACTGGATACGACTCAGTT* | *TGTGCAAATCTATGCAAAACTGA* |
| *mir-19b-3p* | *GTCGTATCCAGTGCAGGGTCCGAGGTATTCGCACTGGATACGACTCAGTT* | *TGTGCAAATCCATGCAAAACTGA* |
| *mir-301a-3p* | *GTCGTATCCAGTGCAGGGTCCGAGGTATTCGCACTGGATACGACGCTTTG* | *CAGTGCAATAGTATTGTCAAAGC* |
| *mir-Reverse* | *ATCCAGTGCAGGGTCCGAGG* |  |

**Supplementary Table S2.** *Primers and stem-loop primers used for miRNAs RT-qPCR experiments.*

| **Antibody** | **Specie** | **Brand** | **N° Catalogue** | **Dilution** |
| --- | --- | --- | --- | --- |
| α-ACTIN | Polyclonal-Goat | Santa Cruz | sc-1616 | 1/1000 |
| α-E2F1 | Monoclonal-Rabbit | Abcam | ab108374 | 1/1000 |
| α-E2F2 | Monoclonal-Mouse | Santa Cruz | sc-9967 | 1/1000 |
| α-E2F3A | Polyclonal-Rabbit | Santa Cruz | sc-879 | 1/1000 |
| α-E2F4 | Monoclonal-Mouse | Santa Cruz | sc-398543 | 1/1000 |
| α-E2F5 | Monoclonal-Mouse | Santa Cruz | sc-374268 | 1/1000 |

**Supplementary Table S3.** *Primary antibodies used for western blot experiments.*

| **miRNA** | **Target genes** | | | | | | | | | |
| --- | --- | --- | --- | --- | --- | --- | --- | --- | --- | --- |
| ***hsa-miR-4517*** | VPS13B | ZNF516 | ADD2 | VPS13B | RCBTB1 | ZRANB2 | ANKRD44 | PRNP | OLA1 | KCNMA1 |
| ***hsa-miR-515-5p*** | JARID2 | DCLK1 | PEX11B | GFRA1 | ECHDC1 | GFRA1 | MRPS33 | TMEM127 | RNF150 | TIMELESS |
| ***hsa-miR-5688*** | NFIA | AHCY | CEP97 | EXOSC6 | ALG14 | VAPA | NIBAN3 | PIK3CD | HP1BP3 | SERBP1 |
| ***hsa-miR-145-5p*** | ABRACL | AAGAB | SEZ6L2 | QRICH1 | UGCG | EHD1 | ZFP37 | GNAI2 | MAP3K9 | PDE1B |
| ***hsa-miR-19a-3p*** | XRN1 | ZZZ3 | FBXO47 | NONO | KCNK3 | RPRM | PRORP | SSBP2 | MS4A2 | ODAPH |
| ***hsa-miR-19b-3p*** | OR1I1 | STK24 | NFATC2 | NTNG1 | TRAK2 | PAK1 | ADAM17 | DIPK1A | HNRNPU | GFRA1 |
| ***hsa-miR-363-5p*** | KCNJ15 | B3GALNT2 | PIK3CA | ZNF718 | C1orf116 | LIN54 | DNAAF10 | FSD1L | RMND5B | ACP3 |
| ***hsa-miR-1323*** | IL21 | NMUR1 | URM1 | AAGAB | DPF3 | ZNF343 | SLC13A5 | TEX30 | SEPTIN9 | FGD4 |
| ***hsa-miR-373-5p*** | HHLA1 | SMG6 | ORAI2 | NISCH | GPR132 | RFX3 | SMG6 | GUCD1 | SLC17A4 | USP7 |
| ***hsa-miR-548ab*** | CLRN1 | SLC9A3 | CDC27 | ALG10 | DCUN1D3 | CLRN1 | DENND4C | SLC9A3 | DENND4C | SIGLEC6 |
| ***hsa-miR-548ah-3p*** | PRKAR1A | BTBD10 | MRPL30 | LYNX1 | PRKAR1A | CYP1B1 | DNASE1L3 | BEST3 | FGF10 | TSKU |
| ***hsa-miR-548av-3p*** | TCEANC2 | NWD1 | GINS1 | NACA | NWD1 | OR4K13 | CYP1B1 | OR4K13 | KIAA0232 | COX20 |
| ***hsa-miR-548o-3p*** | PROX1 | PRKAR1A | MON1B | SYNM | PRKAR1A | TACR1 | TPGS2 | BEST3 | OR2C3 | TPGS2 |
| ***hsa-miR-548p*** | SSH2 | NF2 | PLCB1 | SSH2 | GGA2 | PLCB1 | MTUS1 | MAP2K6 | COL4A5 | HCN1 |
| ***hsa-miR-1272*** | KLHL24 | ZNF287 | GFPT1 | PLEKHA5 | MDM2 | ZNF286A | PEX5 | TMEM59 | SLC6A11 | LSAMP |
| ***hsa-miR-129-2-3p*** | FAM153B | CGGBP1 | MTA2 | DLC1 | DGKB | RPL10 | CDK6 | GNA12 | DUSP22 | TBC1D13 |
| ***hsa-miR-18b-3p*** | DUSP14 | DDX6 | PPP2R2B | IGSF9B | ACO1 | SLC25A17 | PLXDC2 | USP8 | AZIN2 | SLC25A18 |
| ***hsa-miR-516a-5p*** | BMI1 | COMMD3-BMI1 | RCAN1 | SLC2A14 | SRSF8 | FGF13 | IL36RN | SHC4 | TBL2 | ZBTB20 |
| ***hsa-miR-10b-5p*** | CDKN2A | CDKN1A | NR4A3 | ERLIN1 | PTPRT | ORAI2 | SURF6 | TSPAN33 | THUMPD1 | SETD7 |
| ***hsa-miR-186-5p*** | AMD1 | ATL2 | SEPTIN11 | NKAP | ATXN1 | EEF1A1 | FGFR2 | RBM28 | PPP4R2 | 37104 |
| ***hsa-miR-335-5p*** | PIGN | DUSP5 | FOXA2 | ANTXR2 | ATP2B4 | FAM120B | PRPF40B | CHRM2 | LIFR | LZTFL1 |
| ***hsa-miR-96-5p*** | HTR1B | ECT2 | NKIRAS2 | DDIT3 | PHF19 | SGK3 | UBE2N | PPP1R12A | RAB5B | NIPA1 |
| ***hsa-miR-140-5p*** | MMP13 | FGF2 | ARIH1 | FXR1 | SPCS3 | PHACTR2 | RALA | PRDM1 | MYO6 | ZNF426 |
| ***hsa-miR-548ad-5p*** | ANKEF1 | PTBP2 | KATNAL1 | DCUN1D3 | DENND4C | ERCC8 | KCNMA1 | IFT22 | SLMAP | ZNF121 |
| ***hsa-miR-589-3p*** | CDKN1A | CALM3 | CYP26B1 | NPAS3 | LIPG | FGF1 | DOT1L | GLYR1 | VIPR2 | GLYR1 |
| ***hsa-miR-7974*** | SDC4 | NUFIP2 | P3H1 | HMGA1 | FHL2 | PEG10 | PALM2AKAP2 | SMG1 | THRA | KCNIP3 |
| ***hsa-miR-500a-3p*** | SOAT1 | RTF2 | MDM2 | SLC33A1 | TAF8 | OPA3 | PARD6B | GM2A | SOD2 | MIXL1 |
| ***hsa-miR-548ae-5p*** | ANKEF1 | PTBP2 | KATNAL1 | DCUN1D3 | DENND4C | ERCC8 | KCNMA1 | IFT22 | SLMAP | ZNF121 |
| ***hsa-miR-548ay-5p*** | ANKEF1 | PTBP2 | ZNF154 | KATNAL1 | CDC27 | ALG10 | DCUN1D3 | DENND4C | ERCC8 | MCIDAS |
| ***hsa-miR-548d-5p*** | PDE12 | PTCHD1 | ANKEF1 | PTBP2 | ZNF154 | KATNAL1 | PCGF5 | MTURN | BTRC | CDC27 |
| ***hsa-miR-1307-5p*** | PLXNA1 | EDEM3 | IHH | TET3 | SESN3 | CCDC25 | ZFP90 | ZNF618 | SH3BP5L | FAM3B |
| ***hsa-miR-301a-3p*** | KDELR2 | CYBRD1 | PTEN | RAB5B | BTBD3 | TENT2 | TXNIP | NUS1 | SNAPIN | EOGT |
| ***hsa-miR-454-3p*** | PHF12 | UBR7 | PTPRG | TENT2 | LMLN | ATP6V1C1 | DPYSL2 | NFIB | G6PC1 | SERINC3 |
| ***hsa-miR-548av-5p*** | ATXN1L | NRCAM | NRCAM | SLITRK4 | C9orf153 | PTCD2 | PTCD2 | PTCD2 | RPS6KC1 | RPS6KC1 |
| ***hsa-miR-548k*** | MTHFD2 | FEM1C | KATNAL1 | ATXN1L | ARPIN | GRAMD2B | DLGAP1 | TMEM150A | DLGAP1 | C11orf58 |
| ***hsa-miR-1286*** | DCP2 | TNRC6B | RNF38 | SHISA6 | MGRN1 | JADE2 | ATP6AP1 | SNX29 | TEAD1 | ENOX2 |
| ***hsa-miR-1295a*** | TEX261 | TWNK | FAM241A | NOL9 | NUDCD2 | BSND | NT5C1A | MFAP5 | RAB5B | KSR2 |
| ***hsa-miR-21-3p*** | CLIP4 | SREK1IP1 | CYCS | MAT2A | ARNTL | GPAM | KIF6 | DNAJC10 | ARRDC2 | FAM241A |
| ***hsa-miR-33a-5p*** | ABCA1 | CROT | BRWD1 | ZC3H12C | BTBD2 | PPP1R9A | XIAP | HADHB | ABHD2 | UVRAG |
| ***hsa-miR-5701*** | H3-3B | SCML2 | ZC3H7A | PSME3 | USP6NL | NGEF | CEP57L1 | MEGF8 | ZNF468 | PDK1 |
| ***hsa-miR-1322*** | PEDS1 | H6PD | FAM71F2 | MDM2 | C11orf58 | KCNK12 | PARVA | HIF1A | ZNF662 | PPP1R16B |
| ***hsa-miR-33b-5p*** | BCL2 | CREB1 | RMND5A | ABHD2 | ZNF583 | YWHAG | SEMA7A | PLEKHA8 | 37469 | PALLD |
| ***hsa-miR-378d*** | REST | ESYT2 | CACNG8 | TGFB2 | MYRF | LGSN | PPARGC1B | WDR33 | ZNF773 | C3orf70 |
| ***hsa-miR-4286*** | KLHDC3 | TAOK1 | TMEM151B | SEMA4D | SAR1A | PGPEP1 | FCGR3B | SLC5A2 | SCNN1G | EPB41L1 |
| ***hsa-miR-4454*** | ZNF699 | GNE | SLC35F5 | MCTS1 | ACOT9 | RNF4 | KNSTRN | BMF | NFIA | INO80C |
| ***hsa-miR-1260a*** | UNC13A | SLC7A5 | REL | GNPNAT1 | BTG2 | RSL1D1 | KIFC1 | SLC25A42 | AMDHD2 | TNFSF14 |
| ***hsa-miR-1260b*** | TMEM167A | STMN3 | CDKN1B | PHF5A | RPL3 | ZNF233 | EIF4G1 | SLC25A3 | CRK | USP12 |
| ***hsa-miR-1268a*** | CAMK2G | ZNF641 | NOM1 | FAM228A | CERS1 | FRMPD4 | PDE6B | ERO1A | EPB41 | GMPS |
| ***hsa-miR-1268b*** | ZNF641 | TRIM72 | NOM1 | CERS1 | FRMPD4 | PDE6B | EPB41 | RAET1E | OR7D2 | OR7D2 |
| ***hsa-miR-1275*** | IGF1R | ANKRD13B | GDI1 | ARIH1 | MEN1 | SUV39H1 | CAMK2A | UBE2V1 | UBE2N | PEDS1-UBE2V1 |
| ***hsa-miR-498-5p*** | DCUN1D4 | TENT4B | CLDND1 | ZNF765 | SCN2A | NOSTRIN | ZNF705D | USP9X | GK5 | ETNK1 |
| ***hsa-miR-498-3p*** | SLC12A6 | SMIM8 | CARHSP1 | SUMF2 | TGFB1I1 | NSD2 | PSPC1 | CYLD | USP44 | CEP63 |
| ***hsa-miR-1273a*** | no ID match our ID crieria | | | | | | | | | |

**Supplementary Table S4.** Differentially expressed miRNAs (DEGs) and 10 gene targets for each miRNA with the higher *p*-binding.

**Supplementary figures:**

***Analysis of E2F1, E2F2, E2F3A, E2F3B, E2F4 and E2F5 mRNA expression levels in G1-synchronized hPSCs and HF populations***

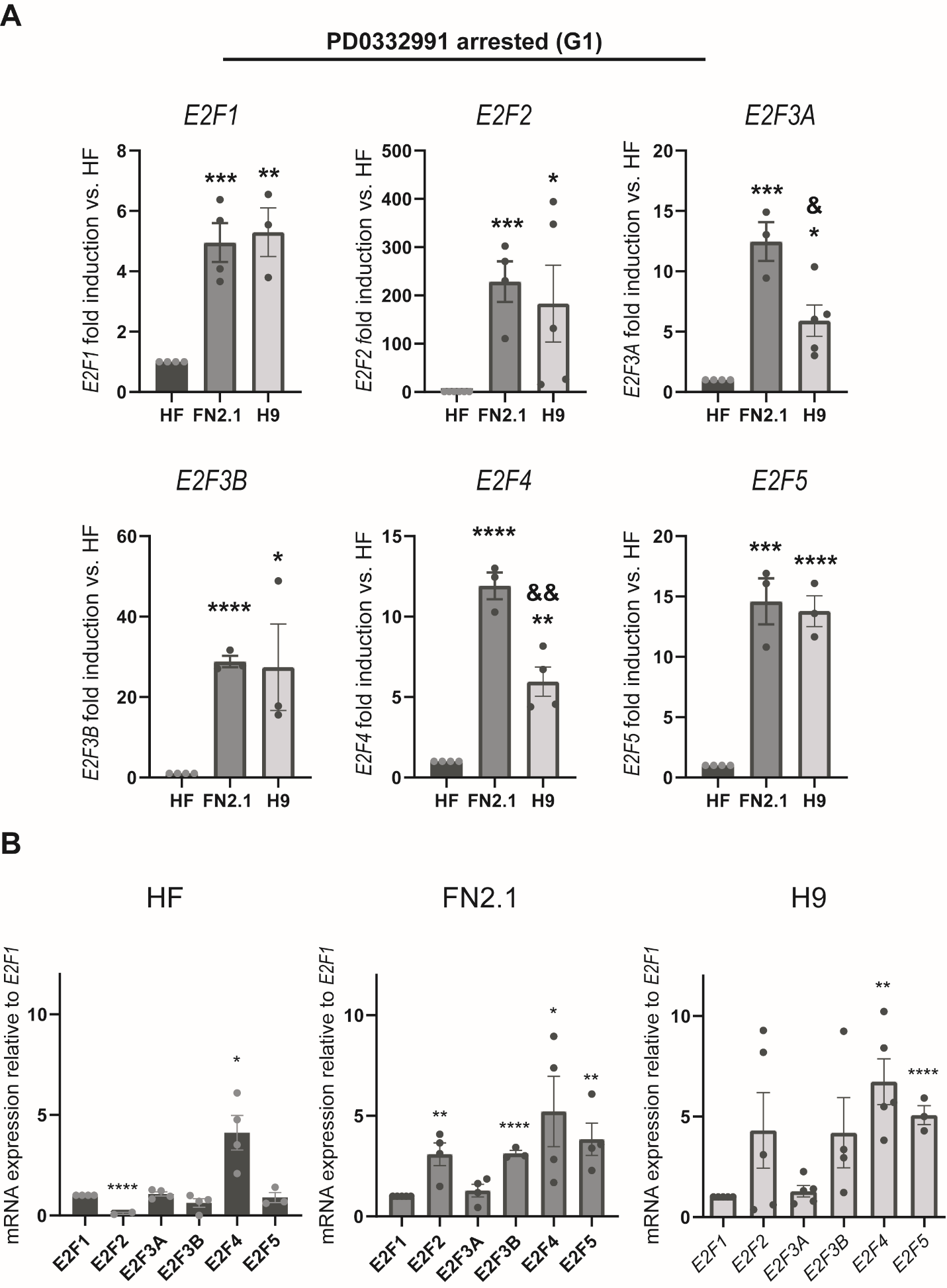

**Supplementary Figure S1. *mRNA expression levels of E2F factors in hPSCs and HF arrested with PD0332991.* (A)** Analysis of *E2F1*, *E2F2*, *E2F3A*, *E2F3B*, *E2F4* and *E2F5* mRNA expression levels, quantified by RT-qPCR, in H9 hESCs and FN2.1 hiPSCs following treatment with PD0332991. *RPL7* expression was used as a normalizer. Graphs show mean ± SEM mRNA fold induction relative to human fibroblast cells (HF) (arbitrarily set as 1) of at least three independent experiments. Statistical analysis was done by Student’s t-test, (*) *p*<0.05; (**) *p*<0.01; (***) *p*<0.001; (****) *p*<0.0001. vs HF (control of differentiated cells). **(B)** Analysis of *E2F1*, *E2F2*, *E2F3A*, *E2F3B*, *E2F4* and *E2F5* mRNA expression levels, quantified by RT-qPCR, within each cell line (HF, FN2.1 hiPSCs and H9 hESCs). *RPL7* expression was used as a normalizer. Graphs show mean ± SEM mRNA fold induction relative to *E2F1* (arbitrarily set as 1) of at least three independent experiments. Statistical analysis was done by Student’s t-test, (*) *p*<0.05; (**) *p*<0.01; (***) *p*<0.001; (****) *p*<0.0001. vs *E2F1* mRNA expression levels within each cell line.

***Cell viability and cell death upon E2F inhibition***

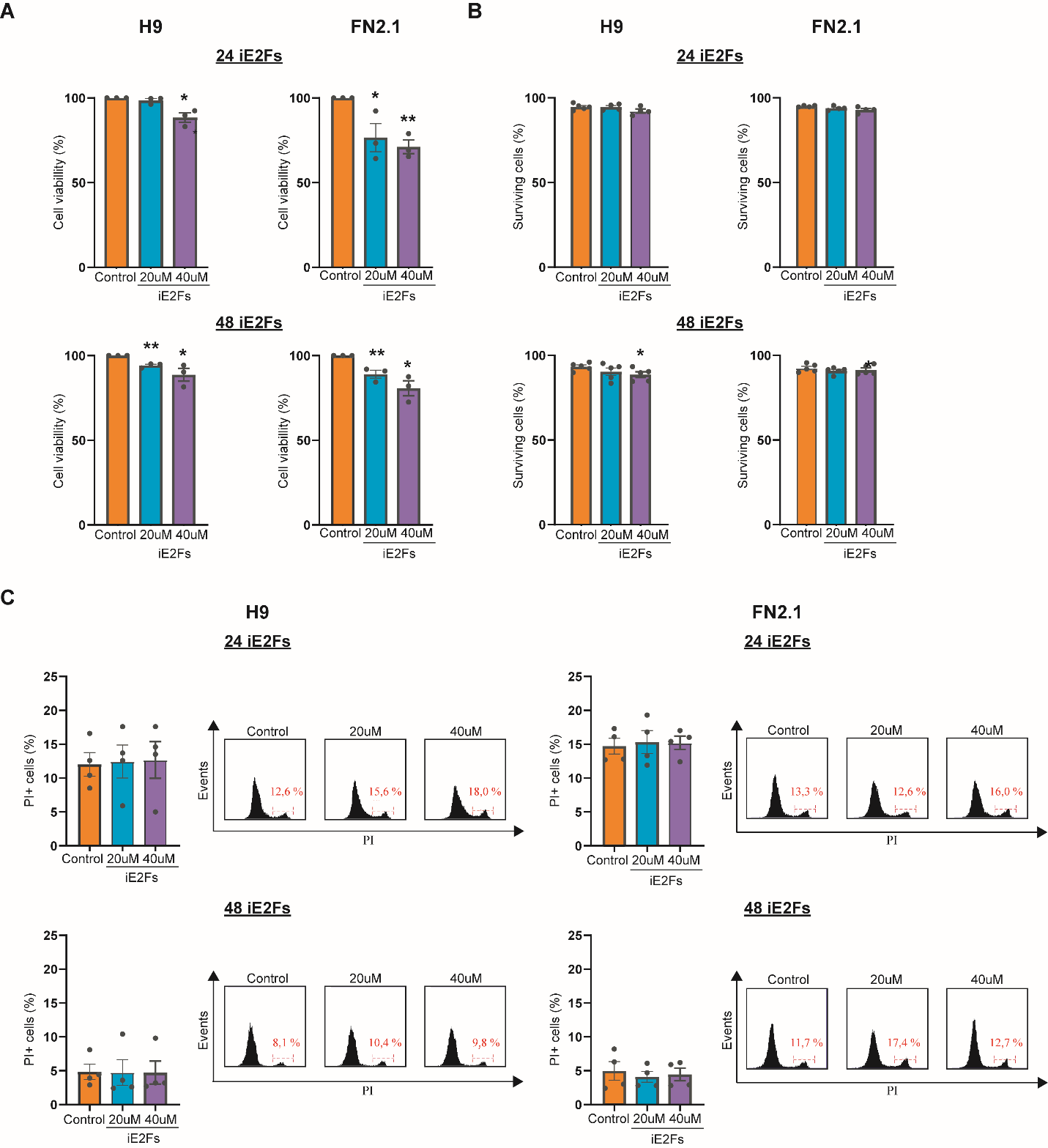

**Supplementary Figure S2. *Changes in cell viability and cell death of hPSCs treated with HLM006474 pan-E2Fs inhibitor.* (A)** H9 and FN2.1 cell viability was analyzed 24 and 48 h post-HLM006474 treatment (20 and 40 μM) by XTT colourimetric assay. Mean ± SEM from three independent experiments are shown. Statistical analysis was done by Student’s t-test, (*) *p*<0.01 and (**) *p*<0.001 vs. Control (DMSO). **(B)** Bar graphs show the percentage of surviving cells assessed by the Trypan blue exclusion method 24 and 48 h after HLM006474 treatment (20 and 40 μM). Mean ± SEM from five independent experiments are shown. Statistical analysis was done by Student’s t-test, (*) *p*<0.01 vs. Control (DMSO). **(C)** Representative histograms of Propidium iodide (PI) stained H9 and FN2.1 unfixed hPSCs treated for 24 and 48 with 20 μM and 40 μM pan-E2F inhibitor. The percentage of PI-positive cells (late apoptotic or necrotic) was determined by flow cytometric analysis. Mean ± SEM from five independent experiments are graphed.

***Bioinformatic analysis and diagnostic plots***

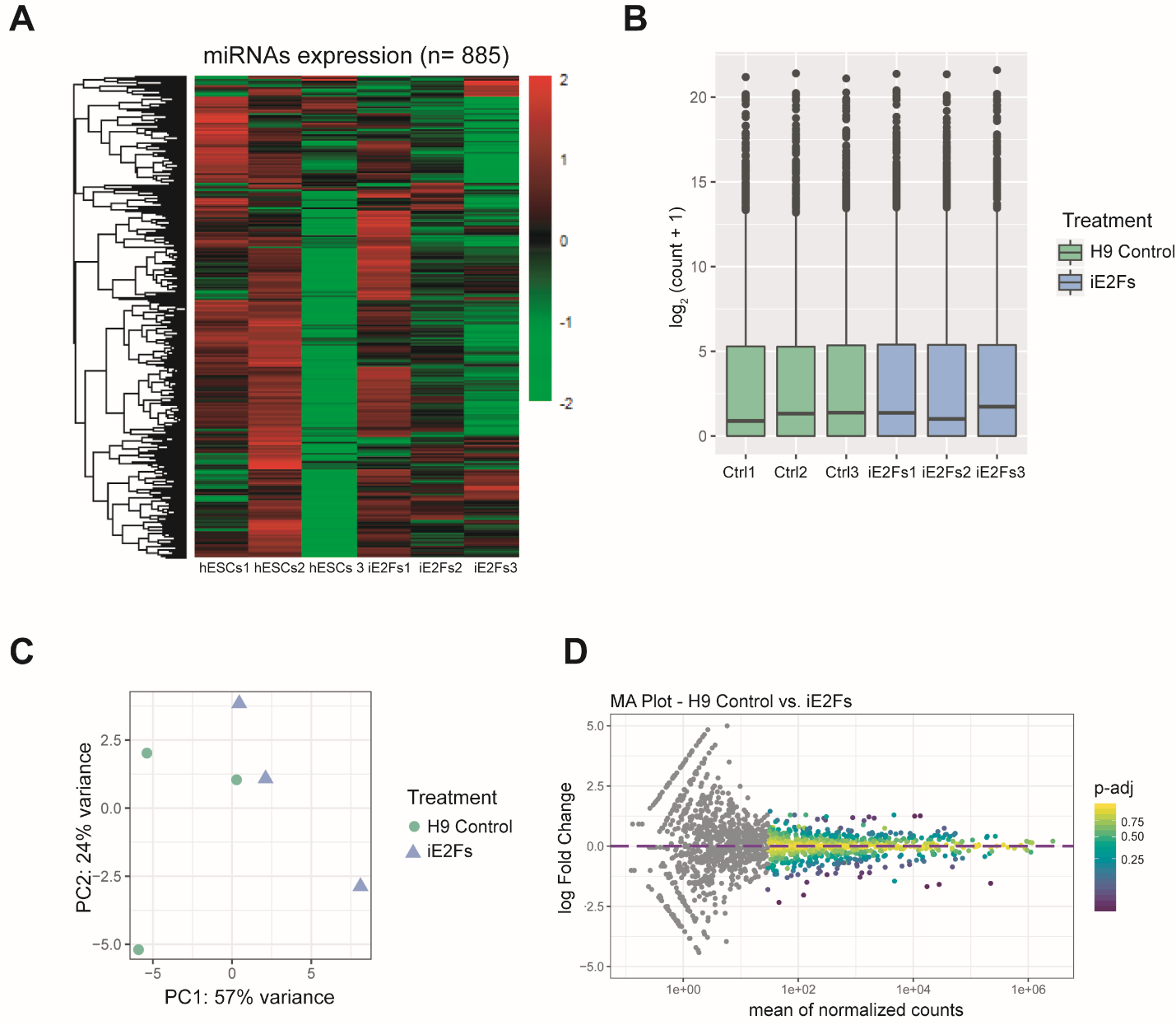

**Supplementary Figure S3. Bioinformatic analysis and diagnostic plots.** **(A)** Heatmap with levels of miRNAs expression of all replicates of control and iE2Fs-treated (24 h, 20 μM) H9 hESCs. miRNAs with readings > 10 were filtered. With the resulting miRNAs, expression levels were transformed to fit on a colour scale between 2 and -2. The heatmap was made with the pheatmap package in R, showing the miRNA expression profile between both samples. **(B)** Box plots represent miRNA expression distribution of the three biological replicates of each cell population: H9 hESCs control and iE2Fs treatment. **(C)** Principal component analysis (PCA) plot shows how replicates of both, H9 hESCs control and iE2Fs treated, are grouped according to the similarity of the data obtained from DESeq2. **(D)** MA-plot compares the log2 fold change of the H9 hESCs control vs. iE2Fs treatment on the y-axis; and the mean of normalized counts by the size factor on the x-axis. A dot represents each miRNA. The colour scale shows miRNAs according to their p-value.
